# A computationally efficient clustering linear combination approach to jointly analyze multiple phenotypes for GWAS

**DOI:** 10.1101/2021.11.22.469509

**Authors:** Meida Wang, Shuanglin Zhang, Qiuying Sha

**Affiliations:** Mathematical Sciences, Michigan Technological University, Houghton, MI, USA

## Abstract

There has been an increasing interest in joint analysis of multiple phenotypes in genome-wide association studies (GWAS) because jointly analyzing multiple phenotypes may increase statistical power to detect genetic variants associated with complex diseases or traits. Recently, many statistical methods have been developed for joint analysis of multiple phenotypes in genetic association studies, including the Clustering Linear Combination (CLC) method. The CLC method works particularly well with phenotypes that have natural groupings, but due to the unknown number of clusters for a given data, the final test statistic of CLC method is the minimum p-value among all p-values of the CLC test statistics obtained from each possible number of clusters. Therefore, a simulation procedure must be used to evaluate the p-value of the final test statistic. This makes the CLC method computationally demanding. We develop a new method called computationally efficient CLC (ceCLC) to test the association between multiple phenotypes and a genetic variant. Instead of using the minimum p-value as the test statistic in the CLC method, ceCLC uses the Cauchy combination test to combine all p-values of the CLC test statistics obtained from each possible number of clusters. The test statistic of ceCLC approximately follows a standard Cauchy distribution, so the p-value can be obtained from the cumulative density function without the need for the simulation procedure. Through extensive simulation studies and application on the COPDGene data, the results demonstrate that the type I error rates of ceCLC are effectively controlled in different simulation settings and ceCLC either outperforms all other methods or has statistical power that is very close to the most powerful method with which it has been compared.

## Introduction

Genome-wide association study (GWAS) has successfully identified a large number of genetic variants that are associated with human complex diseases or phenotypes [1–4]. Among these results, a phenomenon in which a genetic variant affects multiple phenotypes often occurs [5], which is significant evidence to show that pleiotropic effects on human complex diseases are universal [6–9]. Moreover, several disease-related phenotypes are usually measured simultaneously as a disorder or risk factors of a complex disease in GWAS. Therefore, considering the correlated structure of multiple phenotypes in genetic association studies can aggregate multiple effects and increase the statistical power [10–15].

At present, a variety of approaches that focus on jointly analyzing multiple phenotypes have been proposed. These statistical methods can be roughly divided into three categories, including approaches based on regression models [16–19], combining the univariate analysis results [20–23], and variable reduction techniques [24–27]. For example, MultiPhen [19] performs an ordinal regression model, which uses an inverted model whereby the phenotypes are the predictor variables and the genotype is the dependent variable [28–29]. In terms of the second category, combining the univariate test statistics or integrating the p-values of univariate tests are two basic methods. For instance, the O’Brien [20–21] method constructs a test statistic for pleiotropic effect by combining univariate test statistics of multiple phenotypes; the Trait-based Association Test that uses the Extended Simes procedure (TATES) [23] integrates the p-values from univariate tests to obtain an overall trait-based p-value. In addition, principal components analysis of phenotypes (PCP) [24], principal component of heritability (PCH) [25–26], and canonical correlation analysis (CCA) [27] are three variable reduction methods in the third category.

In practice, multiple phenotypes considered may be in different clusters, but most methods for detecting the association between multiple phenotypes and genetic variants either treat all phenotypes as a group or treat each phenotype as one group and combine the results of univariate analysis. Unlike these methods, the clustering linear combination (CLC) method [30] works particularly well with phenotypes that have natural clusters. In the CLC method, individual statistics from the association tests for each phenotype are clustered into positively correlated clusters using the hierarchical clustering method, then the CLC test statistic is used to combine the individual test statistics linearly within each cluster and combine the between-cluster terms in a quadratic form. It was theoretically proved that if the individual statistics can be clustered correctly, the CLC test statistic is the most powerful test among all tests with certain quadratic forms [30]. Due to the unknown number of clusters for a given data, the final test statistic of CLC method is the minimum p-value among all p-values of the CLC test statistics obtained from each possible number of clusters. Therefore, a simulation procedure must be used to evaluate the p-value of the final test statistic because it does not have an asymptotic distribution, and that makes the CLC method computationally demanding. If we can construct a test statistic with an approximate distribution, the computational efficiency will be greatly improved. In this paper, based on the Aggregated Cauchy Association Test (ACAT) method [31], we develop a new method named computationally efficient CLC (ceCLC). In ceCLC, the p-values of the CLC test statistics with *L* clusters are transformed to follow a standard Cauchy distribution, then the transformed p-values are combined linearly with equal treatment to obtain the ceCLC test statistic. This test statistic of ceCLC has an approximately standard Cauchy distribution even though there is a correlated structure between multiple phenotypes [32], so the p-value of the ceCLC test statistic can be calculated based on the cumulative density function of standard Cauchy distribution. We perform extensive simulation studies and apply ceCLC to the COPDGene real dataset. The results show that the ceCLC method has correct type I error rates and either outperforms all other methods or has statistical power that is very close to the most powerful method with which it has been compared.

## Materials and Methods

Assume we consider *N* unrelated individuals with *K* correlated phenotypes, which can be quantitative or qualitative, and each individual has been genotyped at a genetic variant of interest. Let *Y_i_* = (*Y*_*i*1_,…, *Y_iK_*)*^T^* represent *K* correlated phenotypes for the *i*th individual (1 for cases and 0 for controls for a qualitative trait) with *i* = 1,2,…, *N*. Let *G_i_* denote the genotypic score for the *i*th individual at the variant, where *G_i_* ∈ {0, 1, 2} corresponds to the number of minor alleles. We suppose that there are no covariates. If there are *p* covariates *Z*_*i*1_,…, *Z_ip_*, we adjust both genotypes and phenotypes for the covariates [33–34] using linear models *G_i_* = *α*_0_ + *α*_1_*Z*_*i*1_ +… + *α_p_Z_ip_* + *ε_i_* and *Y_ik_* = *α*_0*k*_ + *α*_1*k*_*Z*_*i*1_ +… + *α_pk_Z_ip_* + *τ_ik_*, and use the residuals of the respective linear models to replace the original genotypes and phenotypes.

For each phenotype, we consider the following generalized linear model [35]:

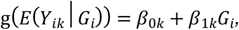

where *β*_1*k*_ is the genetic effect of the variant on the *k*th phenotype and g(·) is a monotone “link” function. Two types of generalized linear model are commonly used: 1) linear model with an identity link for quantitative phenotypes and 2) logistic regression model with a logit link for qualitative phenotypes. We first conduct an univariate test to test *H*_0_: *β*_1*k*_ = 0 for each phenotype, *k* = 1, 2,…, *K*, using the score test statistic [36]

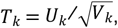

where 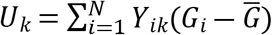 and 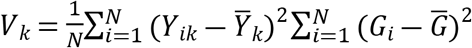. Since the test statistic *T_k_* has an approximate normal distribution with mean *μ_k_* = *E*(*T_k_*) and variance 1, we can assume that *T* = (*T*_1_,…, *T_K_*)*^T^* approximately follows a multivariate normal distribution with mean vector *μ* = (*μ*_1_,…, *μ_K_*)*^T^* and covariance matrix Σ. Our objective is to test the association between multiple phenotypes and a genetic variant, so the null hypothesis is *H*_0_: *β*_11_ =… = *β*_1*K*_ = 0. Sha et al. [30] showed that under the null hypothesis, Σ converges to P(Y) almost surely, where *P*(*Y*) is the correlation matrix of *Y* = (*Y*_1_,…, *Y_K_*)*^T^*. Therefore, we can use the sample correlation matrix of Y, *P^S^*(*Y*), to estimate Σ [30].

Based on the CLC [30] and ACAT methods [31], we propose a computational efficient CLC (ceCLC) method in this paper. Same as the CLC method [30], we use the hierarchical clustering method with similarity matrix 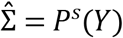 and dissimilarity matrix 1 – *P^S^*(*Y*) to cluster *K* phenotypes. Suppose that the phenotypes are clustered into *L* clusters, considering *L* = 1,…, *K*, and *B* is a *K* × *L* matrix with the (*k,l*)*^th^* element equals 1 if the *k*th phenotype belongs to the /th cluster, otherwise it equals 0. The CLC test statistic [30] with *L* clusters is given by

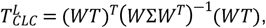

where 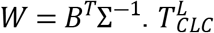 follows a 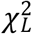 distribution under the null hypothesis, therefore we can obtain the p-value of 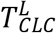, represented by *p_L_*, for *L* = 1,…, *K*. Since for a given data set, the number of clusters of the phenotypes is unknown, in the last step of the CLC method [30], *T_CLC_* = min_1≤*L*≤*K*_ *p_L_* is used as the final test statistic. Because *T_CLC_* does not have an asymptotic distribution, a simulation procedure is needed to evaluate the p-value of *T_CLC_*. This makes the CLC method computationally demanding. In this paper, instead of using the minimum p-value as the test statistic in the CLC method, we use the Cauchy combination test [32] to combine all p-values of the CLC test statistics obtained from each possible number of clusters. We define the ceCLC test statistic as the linear combination of the transformed p-values over the number of *K* clusters, which is given by

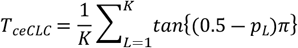

Under the null hypothesis, *tan*{(0.5 – *p_L_*)*π*} follows the standard Cauchy distribution, for *L* = 1,…, *K*. Although there exists a correlated structure between *p*_1_,…, *p_K_*, Liu. et. al [32] has proved that a weighted sum of “correlated” standard Cauchy variables still has an approximately Cauchy tail, and the influence of correlated structure on the tail is quite limited because of the heaviness of the Cauchy tail. Therefore, *T_ceCLc_* can be well approximated by a standard Cauchy distribution. According to the cumulative density distribution of standard Cauchy distribution, the p-value of *T_ceCLc_* can be approximated by 0.5 – {*arctan*(*T_ceCLC_*)/*π*}.

## Results

### Simulation Design

In our simulation studies, we generate one common variant and *K* = 20 and 40 correlated phenotypes for *N* individuals. Firstly, we generate the genotypes of the genetic variant according to the minor allele frequency (MAF = 0.3) under Hardy Weinberg equilibrium. Secondly, the *K* quantitative phenotypes are generated by the following factor model [22, 26, 28, 30]

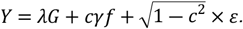

where *Y* = (*Y*_1_,…, *Y_K_*)*^T^, G* is the genotypic score at the variant, *λ* = (*λ*_1_,…, *λ_K_*)*^T^* is the vector of genetic effect sizes on *K* phenotypes, *c* is a constant number, *f* is a vector of factors, and *f* = (*f*_1_,…, *f_R_*)*^T^* ~ *MVN*(0, Σ), where *R* is the number of factors, Σ = (1 – *ρ*) *I* + *ρA*, all elements of matrix *A* equals 1, *I* is an identity matrix, *ρ* is the correlation between factors; *γ* is a *K* × *R* matrix, *ε* = (*ε*_1_,…, *ε_K_*)*^T^* is a vector of residuals, and *ε*_1_,…, *ε_K_* ~ i.i.d. *N*(0, 1).

According to different number of factors affected by the genotypes and different effect sizes, we consider the following four models. In each model, the within-factor correlation is *c*^2^ and the between-factor correlation is *ρc^2^*. We set *c* = 0.5 and *ρ* = 0.6.

Model 1: There is only one factor and genotypes influence all phenotypes. That is, *R* = 1, *λ* = *β*(1, 2,…, *K*)*^T^* and *γ* = (1,…, 1)*^T^*.

Model 2: There are two factors and genotypes influence one factor. That is, *R* = 2, 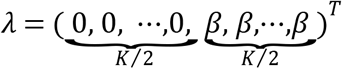, and *γ* = *Bdiag*(*D*_1_, *D*_2_), where *D*_1_ = 1_*K*/2_ for *i*. = 1, 2.

Model 3: There are five factors and genotypes influence two factors. That is, *R* = 5, *λ* = (*β*_11_,…, *β*_1*k*_, *β*_21_,…, *β*_2*k*_, *β*_31_,…, *β*_3*k*_, *β*_41_,…, *β*_4*k*_, *β*_51_,… *β*_5*k*_)*^T^*, and *γ* = *Bdiag*(*D*_1_, *D*_2_, *D*_3_, *D*_4_, *D*_5_), where *D_i_* = 1_*K*/5_ for *i* = 1,…, *k* = *K*/_5_, *β*_11_ =… = *β*_1*k*_ = *β*_21_ =… = *β*_2*k*_ = *β*_31_ =… = *β*_3*k*_ = 0 *β*_41_ =… = *β*_4*k*_ = –*β* and 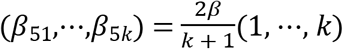.

Model 4: There are five factors and genotypes influence four factors. That is, *R* = 5, *λ* = (*β*_11_,…, *β*_1*k*_, *β*_21_,…, *β*_2*k*_, *β*_31_,…, *β*_3*k*_, *β*_41_,…, *β*_4*k*_, *β*_51_,… *β*_5*k*_)*^T^* and *γ* = *γ* = *Bdiag*(*D*_1_, *D*_2_, *D*_3_, *D*_4_, *D*_5_), where *D_i_* = 1_*K*/5_ for *i* = 1,…, 5, *k* = *K*/5. 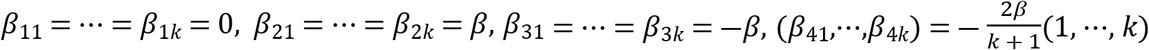, and 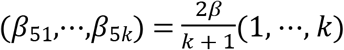.

We consider two types of multiple phenotypes. The first one is that all *K* phenotypes are quantitative and the second one is that half phenotypes are quantitative and the other half are qualitative. To generate a qualitative phenotype, we use a liability threshold model based on a quantitative phenotype. A qualitative phenotype is defined to be affected if the corresponding quantitative phenotype is at least one standard deviation larger (smaller) than the phenotypic mean.

In order to ensure the validity of the ceCLC method, we first evaluate the type I error rates of this method. We simulate data under the null hypothesis and consider three different sample sizes, *N* = 1000, 2000, and 3000, under four different models. The type I error rates are evaluated by 10^5^ replications and at the nominal significance levels of 0.001 and 0.0001, respectively. To evaluate power, we simulate data under the alternative hypothesis and consider two different sample sizes, *N* = 3000 and 5000. The powers are evaluated by 1000 replications at the nominal significance levels of 0.05. To better demonstrate the advantages of the ceCLC method, we compare the performance of ceCLC method with other six existing association tests using multiple phenotypes: CLC [30], MANOVA [37], MultiPhen [19], TATES [23], O’Brien [20], and Omnibus.

### Simulation Results

#### (a) Evaluation of type I error rates

Table 1 presents the type I error rates of the ceCLC method for *K* = 20 quantitative phenotypes, and the type I error rates of the other six methods (CLC, MANOVA, MultiPhen, TATES, O’Brien, Omnibus) are summarized in S1 Table. The corresponding type I error rates for the case of half quantitative traits and half qualitative phenotypes are recorded in Table 2 and S2 Table. In addition, The type I error rate of the ceCLC method for *K* = 40 are listed in S3 and S4 Tables, and the type I error rates of the other six methods for *K* = 40 are summarized in S5 and S6 Tables. For 10^5^ replications, the 95% confidence intervals of Type I error rates divided by nominal significance levels of 0.001 and 0.0001 are (0.8041, 1.1959) and (0.3802, 1.6198), respectively. In the tables, we use gray color to indicate that the type I error rate is not in the 95% CI.

**Table 1.**
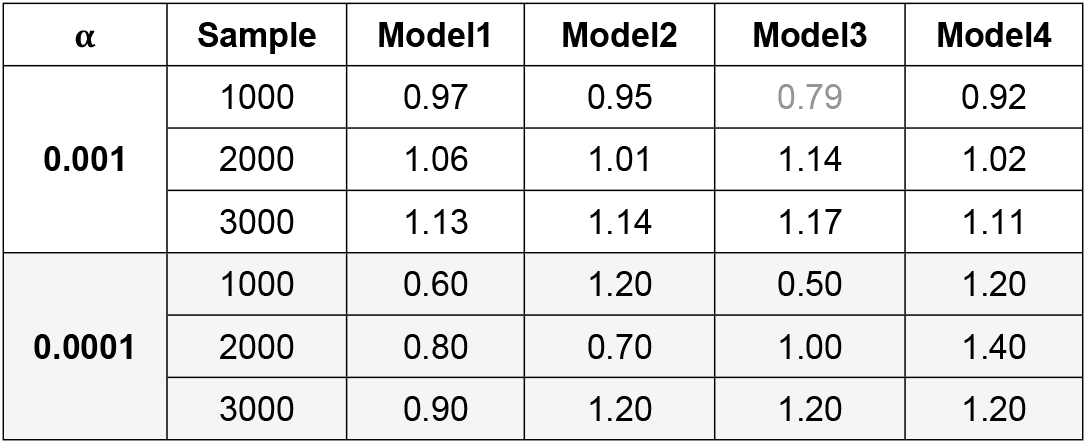
The estimated type I error rates divided by the nominal significance levels of the ceCLC method for 20 quantitative phenotypes.

| $\alpha$ | Sample | Model1 | Model2 | Model3 | Model4 |
| --- | --- | --- | --- | --- | --- |
| **0.001** | 1000 | 0.97 | 0.95 | 0.79 | 0.92 |
|  | 2000 | 1.06 | 1.01 | 1.14 | 1.02 |
|  | 3000 | 1.13 | 1.14 | 1.17 | 1.11 |
| **0.0001** | 1000 | 0.60 | 1.20 | 0.50 | 1.20 |
|  | 2000 | 0.80 | 0.70 | 1.00 | 1.40 |
|  | 3000 | 0.90 | 1.20 | 1.20 | 1.20 |

**Table 2.** The estimated type I error rates divided by the nominal significance levels of the ceCLC method for 10 quantitative and 10 qualitative phenotypes.

| $\alpha$ | Sample | Model1 | Model2 | Model3 | Model4 |
| --- | --- | --- | --- | --- | --- |
| **0.001** | 1000 | 0.98 | 0.93 | 0.87 | 0.91 |
|  | 2000 | 1.00 | 1.09 | 1.06 | 1.03 |
|  | 3000 | 0.97 | 1.08 | 1.02 | 1.04 |
| **0.0001** | 1000 | 0.90 | 1.10 | 1.10 | 1.00 |
|  | 2000 | 0.90 | 0.60 | 1.20 | 1.30 |
|  | 3000 | 0.60 | 0.80 | 1.10 | 1.20 |

From Tables 1 and 2 (S3 and S4 Tables), we can see that all the estimated type I error rates are within the corresponding 95% CIs except 0.79 in Table 1 and 0.76 in S3 Table. However, these two values are very close to the lower bound of its CI (0.8041), therefore we can conclude that the ceCLC method is a valid test. From S1-S2 and S5-S6 Tables, we observe that CLC, MANOVA, TATES, and O’Brien can control type I error rates well, but some of the type I error rates of MultiPhen are slightly inflated and some of the type I error rates of Omnibus are conservative.

#### (b) Assessment of powers

Fig 1 shows the results of power comparisons for all seven tests with 20 quantitative phenotypes when the sample size is 5000. From Fig 1, we find that 1) the ceCLC and CLC methods are more powerful than other methods; 2) the O’Brien method is very sensitive to the direction of the genetic effect on the phenotypes. Its power will decrease dramatically with different directions of the genetic effect on the phenotypes (Models 3 and 4); 3) MANOVA, Omnibus, and MultiPhen have similar powers. Fig 2 shows the results of power comparisons for all seven tests with 10 quantitative and 10 qualitative phenotypes when the sample size is 5000. The general trend of Fig 2 is similar to Fig 1, but the powers of MANOVA, Omnibus, and MultiPhen are higher than those in Fig 1 for Models 3 and 4. S1 and S2 Figs present the results of power comparisons with 40 phenotypes for the sample size of 5000, and all the results of power comparisons for the sample size of 3000 are showed in S3-S6 Figs. In summary, ceCLC and CLC methods are more powerful than other methods under most scenarios ceCLC is much more computationally efficient than CLC.

**Fig 1.**
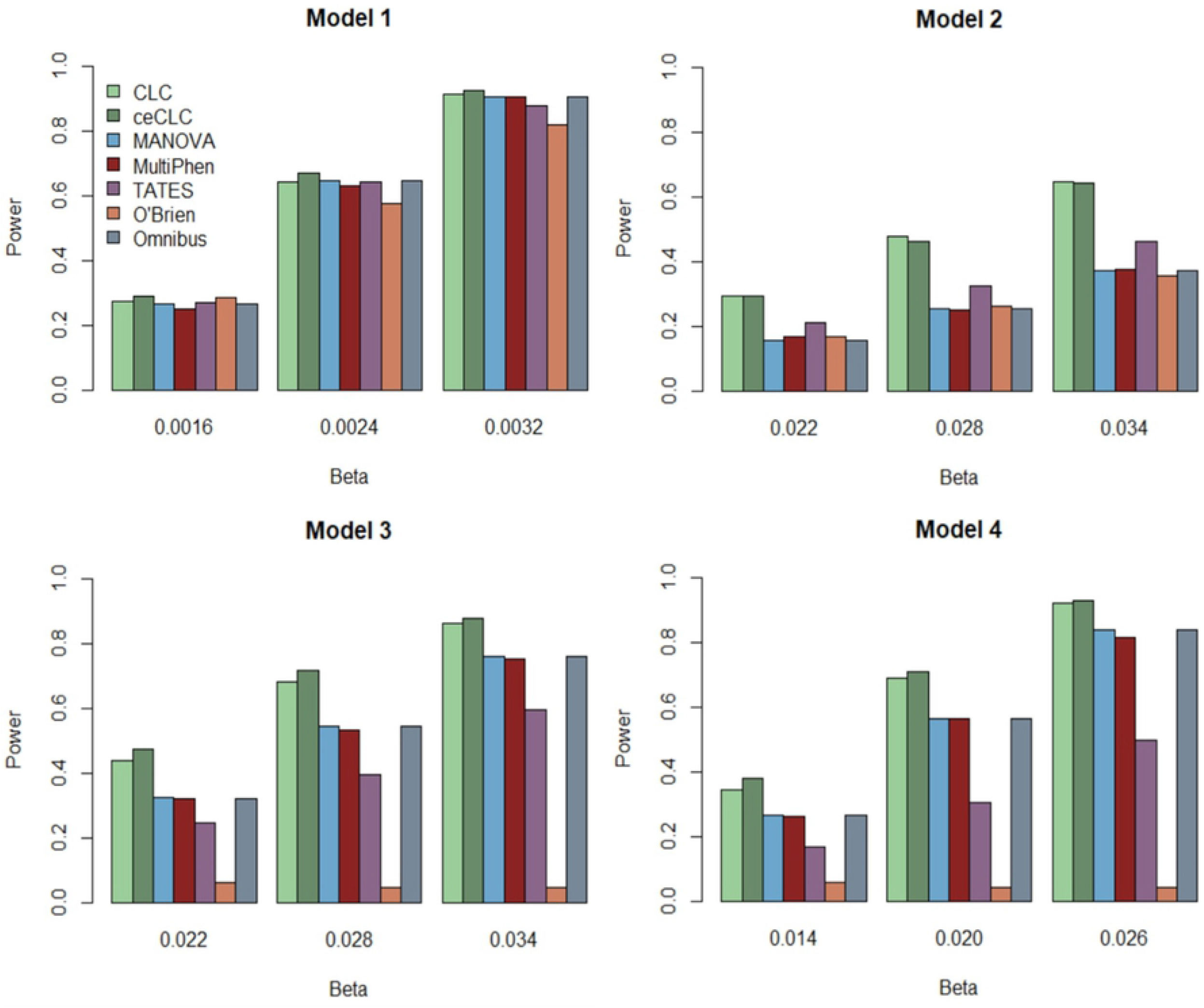
Power comparisons of the seven tests, CLC, ceCLC, MANOVA, MultiPhen, TATES, O’Brien, and Omnibus, with 20 quantitative phenotypes for the sample size of 5000.

**Fig 2.**
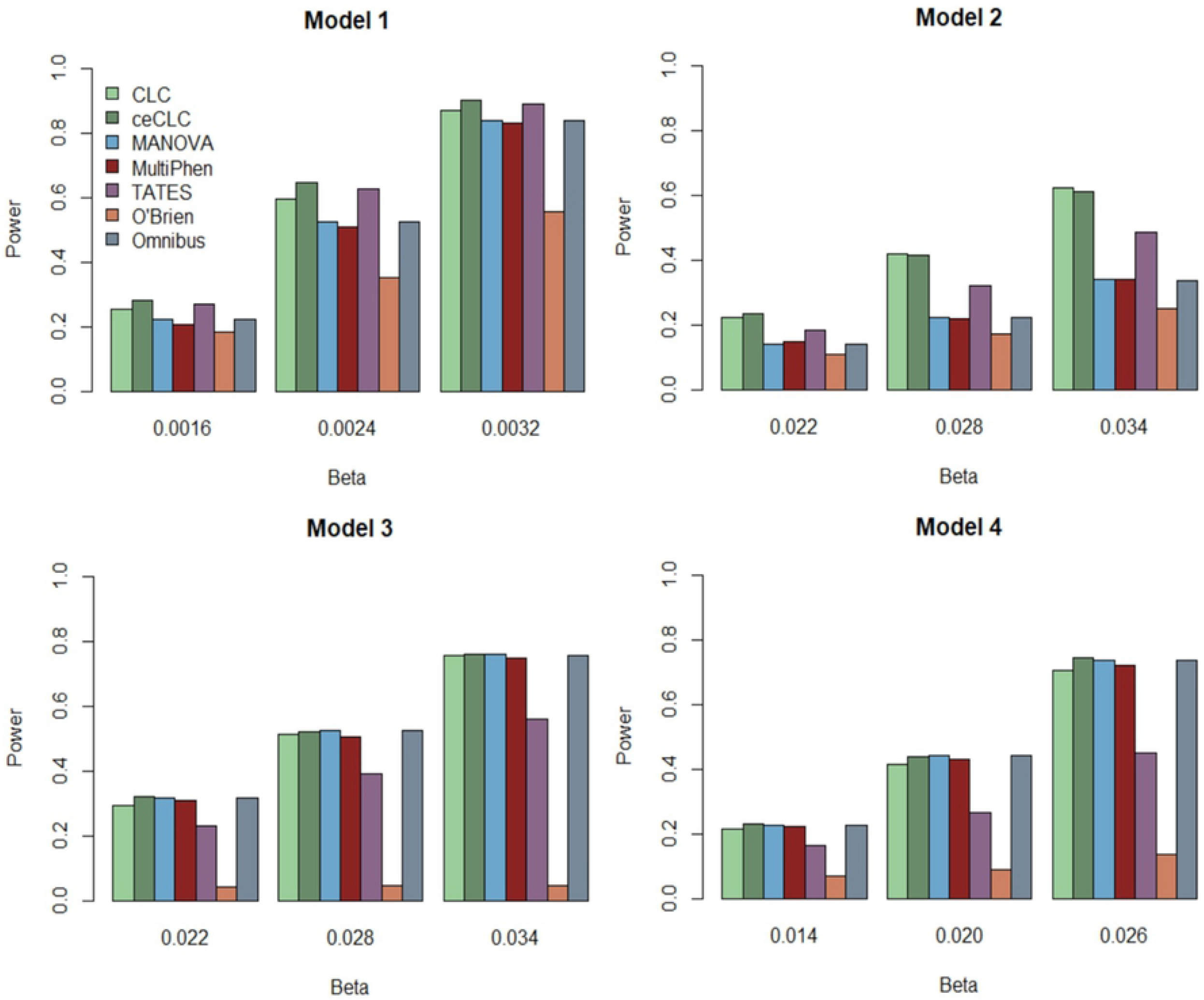
Power comparisons of the seven tests, CLC, ceCLC, MANOVA, MultiPhen, TATES, O’Brien, and Omnibus, with 10 quantitative and 10 qualitative phenotypes for the sample size of 5000.

## Application to the COPDGene Study

Chronic obstructive pulmonary disease (COPD) is a common disease characterized by the presence of expiratory dyspnea due to the excessive inflammatory reaction of harmful gases and particles [38–40]. COPD causes a high mortality and has been reported to be potentially affected by genetic factors [41]. The COPDGene study is a representative multicenter research to detect hereditary factors of this disease [42]. The corresponding dataset of this study has been introduced in our previous papers [22, 30].

Same with the analysis in [22, 30], we consider seven quantitative COPD-related phenotypes, containing FEV1, Emphysema, Emphysema Distribution, Gas Trapping, Airway Wall Area, Exacerbation frequency, and Six-minute walk distance. We also consider four covariates which include BMI, Age, Pack-Years and Sex. After deleting the missing data, there are 5,430 subjects across 630,860 SNPs left for the analysis. We change the sign of six-minute walk distance and FEV1, so that the correlations between the 7 phenotypes are all positive. We apply all seven tests to the processed COPD data.

In our analysis, we choose the commonly used genome-wide significant level *α* = 5 × 10^−8^, Table 3 presents 14 SNPs that are detected by at least one method. All of these 14 SNPs have been reported to be associated with COPD before [43–46]. From Table 3, we can see that MultiPhen detected 14 SNPs; ceCLC, CLC, MANOVA and Omnibus detected 13 SNPs; TATES detected 9 SNPs; and O’Brien only detected 5 SNPs.

**Table 3.** Significant SNPs and the corresponding p-values in the analysis of COPDGene study.

| Chr | Position | Variant identifier | CLC | ceCLC | MANOVA | MultiPhen | TATES | O'Brien | Omnibus |
| --- | --- | --- | --- | --- | --- | --- | --- | --- | --- |
| 4 | 14543149 | rs1512282 | $10^{-9}$ | $5.70 \times 10^{-11}$ | $1.69 \times 10^{-9}$ | $1.03 \times 10^{-9}$ | $5.77 \times 10^{-9}$ | $7.69 \times 10^{-9}$ | $1.82 \times 10^{-9}$ |
| 4 | 14543474 | rs1032297 | $10^{-9}$ | $2.39 \times 10^{-15}$ | $6.52 \times 10^{-14}$ | $7.69 \times 10^{-14}$ | $6.22 \times 10^{-13}$ | $3.35 \times 10^{-10}$ | $7.73 \times 10^{-14}$ |
| 4 | 14547447 | rs1489759 | $10^{-9}$ | 0 | $1.11 \times 10^{-16}$ | $1.22 \times 10^{-16}$ | $2.52 \times 10^{-16}$ | $2.61 \times 10^{-11}$ | $1.11 \times 10^{-16}$ |
| 4 | 14548573 | rs1980057 | $10^{-9}$ | 0 | $6.68 \times 10^{-17}$ | $8.14 \times 10^{-17}$ | $9.35 \times 10^{-17}$ | $3.04 \times 10^{-11}$ | $1.11 \times 10^{-16}$ |
| 4 | 14548591 | rs7655625 | $10^{-9}$ | 0 | $7.12 \times 10^{-17}$ | $9.13 \times 10^{-17}$ | $1.64 \times 10^{-16}$ | $3.08 \times 10^{-11}$ | $1.11 \times 10^{-16}$ |
| 15 | 78882925 | rs16969968 | $10^{-9}$ | $4.91 \times 10^{-11}$ | $1.32 \times 10^{-11}$ | $7.84 \times 10^{-12}$ | $2.98 \times 10^{-8}$ | $9.75 \times 10^{-6}$ | $1.26 \times 10^{-11}$ |
| 15 | 78894339 | rs1051730 | $10^{-9}$ | $4.74 \times 10^{-11}$ | $1.41 \times 10^{-11}$ | $8.16 \times 10^{-12}$ | $2.63 \times 10^{-8}$ | $8.99 \times 10^{-6}$ | $1.35 \times 10^{-11}$ |
| 15 | 78898723 | rs12914385 | $10^{-9}$ | $2.57 \times 10^{-12}$ | $1.76 \times 10^{-12}$ | $1.48 \times 10^{-12}$ | $5.14 \times 10^{-10}$ | $6.12 \times 10^{-8}$ | $1.66 \times 10^{-12}$ |
| 15 | 78911181 | rs8040868 | $10^{-9}$ | $5.08 \times 10^{-12}$ | $2.74 \times 10^{-12}$ | $2.59 \times 10^{-12}$ | $2.40 \times 10^{-9}$ | $1.53 \times 10^{-7}$ | $2.50 \times 10^{-16}$ |
| 15 | 78878541 | rs951266 | $10^{-9}$ | $7.03 \times 10^{-11}$ | $1.77 \times 10^{-11}$ | $1.02 \times 10^{-11}$ | $5.17 \times 10^{-8}$ | $1.50 \times 10^{-5}$ | $1.69 \times 10^{-11}$ |
| 15 | 78806023 | rs8034191 | $10^{-9}$ | $8.03 \times 10^{-10}$ | $2.14 \times 10^{-10}$ | $7.74 \times 10^{-11}$ | $1.02 \times 10^{-7}$ | $2.13 \times 10^{-5}$ | $1.99 \times 10^{-10}$ |
| 15 | 78851615 | rs2036527 | $8.33 \times 10^{-10}$ | $1.52 \times 10^{-9}$ | $3.99 \times 10^{-10}$ | $1.77 \times 10^{-10}$ | $1.56 \times 10^{-7}$ | $2.65 \times 10^{-5}$ | $3.76 \times 10^{-10}$ |
| 15 | 78826180 | rs931794 | $10^{-9}$ | $1.18 \times 10^{-9}$ | $2.35 \times 10^{-10}$ | $9.09 \times 10^{-11}$ | $1.18 \times 10^{-7}$ | $2.33 \times 10^{-5}$ | $2.19 \times 10^{-10}$ |
| 15 | 78740964 | rs2568494 | $3.98 \times 10^{-7}$ | $5.02 \times 10^{-7}$ | $1.05 \times 10^{-7}$ | $4.23 \times 10^{-8}$ | $2.88 \times 10^{-5}$ | $2.38 \times 10^{-3}$ | $9.73 \times 10^{-8}$ |

The p-values of CLC are evaluated using $10^9$ simulations. The p-values of ceCLC, O'Brien, Omnibus, TATES, MANOVA, and MultiPhen are evaluated using their asymptotic distributions. The graying out p-values indicate the p-values $> 5 \times 10^{-8}$ .

## Discussion

In the medical field, many human complex diseases are often accompanied by multiple correlated phenotypes which are usually measured simultaneously, and pleiotropy is a universal phenomenon, so jointly analyzing multiple phenotypes in genetic association studies will very likely increase the statistical power to identify genetic variants that are associated with complex diseases. In this paper, based on the existing CLC method [30] and ACAT [31] strategy, we develop the ceCLC method to test association between multiple phenotypes and a genetic variant. We perform a variety of simulation studies, as well as an application to the COPDGene study to evaluate our new method. The results suggest that the ceCLC method not only has the advantages of the CLC method but is also computationally efficient. The test statistic of the ceCLC method can be well approximated by a standard Cauchy distribution, so the p-value can be obtained from the cumulative density function without the need for the simulation procedure. Therefore, the ceCLC method is computational efficient.

In recent phenome-wide association studies (PheWAS), a great number of phenotypes are collected, which requires powerful and efficient statistical methods. The ceCLC method developed here can be applied to PheWAS. One limitation of the ceCLC method is that it requires individual-level phenotype data and GWAS summary statistics. We know that individual-level phenotype data are often not easily accessible as a result of privacy concerns. Therefore, we are currently considering a new strategy to extend the ceCLC method so that it can be applied to GWAS summary statistics without the requirement for individual-level phenotype data.

## Acknowledgements

This research used data generated by the COPDGene study (phs000179/HMB and phs000179/DS-CS-RD), which was supported by National Institutes of Health (NIH) grants U01HL089856 and U01HL089897. The content is solely the responsibility of the authors and does not necessarily represent the official views of the National Heart, Lung, and Blood Institute or the National Institutes of Health. The COPDGene project is also supported by the COPD Foundation through contributions made by an Industry Advisory Board comprised of Pfizer, AstraZeneca, Boehringer Ingelheim, Novartis, and Sunovion.

## Supporting information

**S1 Table. The estimated type I error rates divided by nominal significance levels of the other six methods, CLC, MANOVA, MultiPhen, TATES, O’Brien, and Omnibus, for 20 quantitative phenotypes.**

**S2 Table. The estimated type I error rates divided by nominal significance levels of the other six methods, CLC, MANOVA, MultiPhen, TATES, O’Brien, and Omnibus for 10 quantitative and 10 qualitative phenotypes.**

**S3 Table. The estimated type I error rates divided by the nominal significance levels of the ceCLC method for 40 quantitative phenotypes.**

**S4 Table. The estimated type I error rates divided by the nominal significance levels of the ceCLC method for 20 quantitative and 20 qualitative phenotypes.**

**S5 Table. The estimated type I error rates divided by nominal significance levels of the other six methods, CLC, MANOVA, MultiPhen, TATES, O’Brien, and Omnibus, for 40 quantitative phenotypes.**

**S6 Table. The estimated type I error rates divided by nominal significance levels of the other six methods, CLC, MANOVA, MultiPhen, TATES, O’Brien, and Omnibus for 20 quantitative and 20 qualitative phenotypes.**

**S1 Fig. Power comparisons of the seven tests, CLC, ceCLC, MANOVA, MultiPhen, TATES, O’Brien, and Omnibus, with 40 quantitative phenotypes for the sample size of 5000.**

**S2 Fig. Power comparisons of the seven tests, CLC, ceCLC, MANOVA, MultiPhen, TATES, O’Brien, and Omnibus, with 20 quantitative and 20 qualitative phenotypes for the sample size of 5000.**

**S3 Fig. Power comparisons of the seven tests, CLC, ceCLC, MANOVA, MultiPhen, TATES, O’Brien, and Omnibus, with 20 quantitative phenotypes for the sample size of 3000.**

**S4 Fig. Power comparisons of the seven tests, CLC, ceCLC, MANOVA, MultiPhen, TATES, O’Brien, and Omnibus, with 10 quantitative and 10 qualitative phenotypes for the sample size of 3000.**

**S5 Fig. Power comparisons of the seven tests, CLC, ceCLC, MANOVA, MultiPhen, TATES, O’Brien, and Omnibus, with 40 quantitative phenotypes for the sample size of 3000.**

**S6 Fig. Power comparisons of the seven tests, CLC, ceCLC, MANOVA, MultiPhen, TATES, O’Brien, and Omnibus, with 20 quantitative and 20 qualitative phenotypes for the sample size of 3000.**

## Notes

### Competing Interest Statement

The authors have declared no competing interest.

